# Identification of five patterns of nucleosome positioning that globally describe transcription factor function

**DOI:** 10.1101/003483

**Authors:** Kazumitsu Maehara, Yasuyuki Ohkawa

## Abstract

Following the binding of transcription factors (TF) to specific regions, chromatin remodeling including alterations in nucleosome positioning (NP) occurs. These changes in NP cause selective gene expression to determine cell function. However whether specific NP patterns upon TF binding determine the transcriptional regulation such as gene activation or suppression is unclear. Here we identified five patterns of NP around TF binding sites (TFBSs) using fixed MNase-Seq analysis. The most frequently observed NP pattern described the transcription state. The five patterns explained approximately 80% of the whole NP pattern on the genome in mouse C2C12 cells. We further performed ChIP-Seq using the input obtained from the fixed MNase-Seq. The result showed that a single trial of ChIP-Seq could visualize the NP patterns around the TFBS and predict the function of the transcriptional regulation at the same time. These findings indicate that NP can directly predict the function of TFs.

## INTRODUCTION

The chromatin structure consists of nucleosomes on genomic DNA. The positions of the nucleosomes (nucleosome positioning; NP) are dynamically determined by transcription factors (TFs) binding to specific DNA sequences. NP alterations by chromatin remodeling are essential for transcriptional activation upon TF binding. Accordingly, NP has been shown to reflect the transcription state directly (Jiang and Pugh 2009; He et al. 2010), and specific NP results from the transcriptional activation or suppression of a gene.

Micrococcal nuclease (MNase) analysis has been widely used to map individual nucleosomes in a locus on the genome (Wu 1980). Recent developments in deep sequencing technology (MNase-Seq) have enabled high-resolution NP analysis (Albert et al. 2007; Mavrich et al. 2008a; Zhang and Pugh 2011), especially in studies of budding yeast. These studies have revealed NP follows specific patterns, and nucleosome depleted region (NDR) (Yuan et al. 2005; Mavrich et al. 2008b), which is considered to be facilitate RNA Polymerase II (RNAP2) recruitment, has been frequently observed by MNase-Seq (Hughes et al. 2012). Albert et al. revealed that the TATA box and transcription start sites (TSS) reside at both ends of the linker DNA in yeast (Albert et al. 2007), suggesting NP determines the start point of transcription. Another characteristic of NP revealed by MNase-Seq is that nucleosomes are aligned at strict and regular intervals as shown in the binding of CTCF, an insulator binding protein that forms boundaries in the genome (Bell et al. 1999; Fu et al. 2008). Recently, Ranjan et al. showed yeast SWR1, a chromatin remodelling enzyme, preferentially recognizes long nucleosome-free DNA (Ranjan et al. 2013), indicating it is a decoder if specific NPs were considered as one of structural code on chromatin that indicates transcription states.

NP has been suggested to be critical for transcription regulation in mammalian genomes due to the absence of the core promoter sequences seen in yeast (Li et al. 2007). The acquisition of high-resolution NP from mammalian genomes is severely limited, because mammal genomes are approximately 100 times larger than yeast’s. Teif et al. used a signal averaging method, average profiling, to demonstrate that nucleosome occupancies could change around lineage specific TFBSs during the differentiation of mouse embryonic stem cells (Teif et al. 2012). These data suggest that the diversity of NP patterns could be formed by lineage-specific TFs.

To clarify NP patterns in certain transcription states in the mammalian whole genome, we developed a novel NP evaluation framework that acquires profiles of the average nucleosome density (PAND) and combined it with fixed MNase-Seq to extract common NP patterns on the genome. Our framework obtained precise NP without a large number of sequencing reads. Using mouse C2C12 myoblast cells, we extracted five common patterns that describe NP around TFBSs on the whole genome. The most frequently observed NP pattern in the five could explain whether transcription was in the active or inactive state. We further demonstrated that PAND is useful for predicting TF function in transcription regulation by a single trial of ChIP-Seq that incorporated fixed MNase-Seq.

## RESULTS

### Fixed MNase-Seq refines NP extraction

To extract NP patterns related to transcription activation or suppression on the whole genome, we attempted to extract NP by MNase analysis on a deep sequencing platform using ∼10^6^ C2C12 cells as a model of the transcription state change in skeletal muscle differentiation despite MNase-Seq normally requiring ∼10^8^ cells to obtain significant NP data. We prepared cross-linked chromatin fractions before MNase digestion to obtain precise NP information. DNA fragments were completely digested into mono-nucleosome size (Fig. 1b, 164 ± 12 bp; mean ± 1 standard-deviation), because the NP data are sensitive to fragment size (Henikoff et al. 2011). The number of mapped reads we acquired was approximately 20 million (myoblast, 21,729,582; myotube, 19,877,072). As expected, the signal was dispersed (Supplementary Fig. 1a); e.g., assuming 20 million reads, the probability that at least one read was detected within a 200 bp interval is approximately 77% if assuming a binomial distribution, a mouse genome size of 2.7 Gbp and nucleosome repeat length of 200 bp.

**Figure 1.**
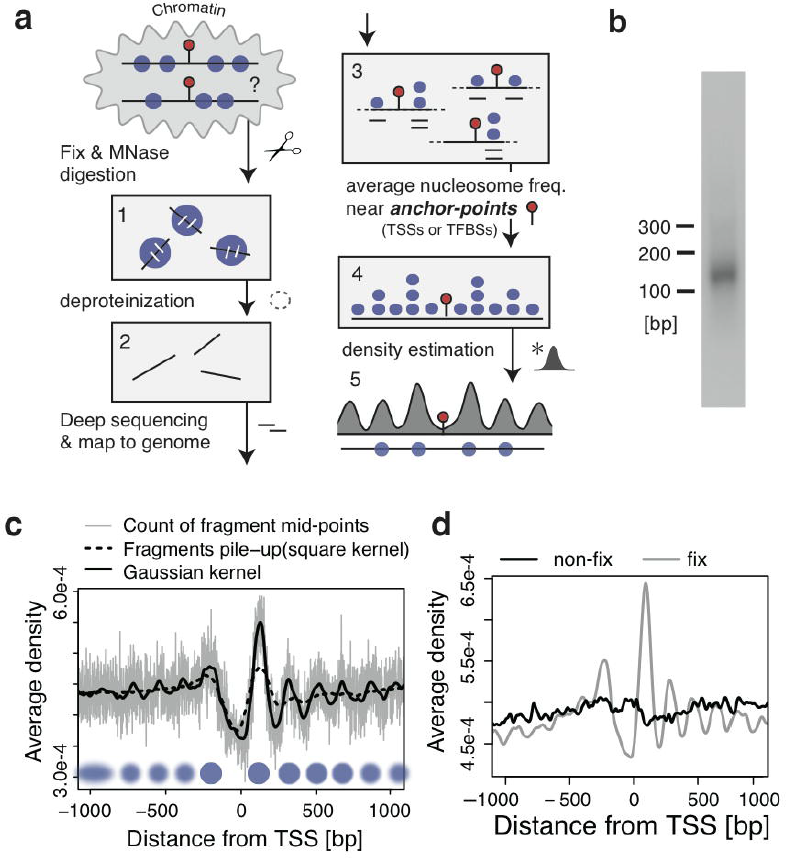
Density estimation of nucleosomes from fixed MNase-Seq. a. **(a)** Analysis flow of average nucleosome density estimation. First, chromatin are fixed and digested by MNase to obtain mono-nucleosome sizes of DNA (1). The samples are deproteinized (2), deep sequenced and map the DNA onto the reference genome (3). Next, the nucleosome detection frequency is averaged at all defined anchor-points (e.g. TSSs, TFBSs) (4). Finally, NP is predicted by the smoothed density profile of the nucleosomes (5).
b. **(b)** DNA fragment size of the ChIP-Seq (RNAP2-S5ph) samples used. DNA fragments of mono-nucleosome size (164+/−12bp) were obtained.
c. **(c)** Comparison of signal processing methods for single PAND analysis. The X-axis shows the distance (in bp) from the TSSs, and the Y-axis shows nucleosome density averaged over all genes. The grey line represents counts of fragment mid-points with 1 bp resolution. The black solid line shows these counts smoothed by a Gaussian kernel, and the dotted line represents the result of fragments pile-up in which mapped reads are extended to the fragment length and accumulated. The blue circles represent inferred positions of the nucleosome (blur: fuzzy, solid: well-positioned).
d. **(d)** A PAND within 1 kbp of a TSS in fixed (grey) and non-fixed (black) samples. The effect of fixation was evaluated for mono-nucleosome sized samples. See Supplementary Figure 1d for fragment size. Fixed samples show clearer positioning around TSSs.

To overcome the signal dispersion for detecting NP, we applied the average profiling method, which is commonly used to evaluate signal distributions in ChIP/MNase-Seq studies (Teif et al. 2012; Wang et al. 2012; Maehara et al. 2013). However, NP could not be clearly captured with this method alone (Fig. 1c, gray, dotted black line). We therefore combined it with the density estimation method to acquire PANDs and precise estimates of NP that are consistent with NDR having +/−1 nucleosomes flanking the TSS (Fig. 1c, solid black line). A similar result of PANDs was also obtained from the input-DNA control data of ChIP-Seq by Asp et al., which used fixed chromatin digested with MNase treatment (Supplementary Fig. 1b) (Asp et al. 2011). We further evaluated PANDs around CTCF binding sites and confirmed stable NP as previously reported (Supplementary Fig. 1c) (Fu et al. 2008). We also confirmed that fixation is preferred for NP detection (Fig. 1d and Supplementary Fig. 1d).

### Five common NP patterns in C2C12 cells

The PAND at TSSs and CTCF binding sites extracted not only NP patterns, but also showed the capability of exploring not-yet characterised NP. Therefore we investigated variable NP using a *cis*-element database from TRANSFAC (Matys et al. 2006). Each *cis*-element resulted in 258 PANDs from fixed MNase-Seq data. Representative data of the MyoD *cis*-element (MYOD_Q6) showed increased nucleosome occupancy around its binding sequence (Fig. 2a). In contrast, the Oct1 *cis*-element (OCT1_07) had decreased nucleosome occupancy near its binding sequence (Fig. 2b). Each of the 258 PANDs had unique shapes (Supplementary Fig. 2). In some PANDs, sharp dips and peaks were observed around +/−100 bp from the center of the *cis*-element, which could be explained by the sequence specificity of MNase (Fig. 2a, b).

**Figure 2.**
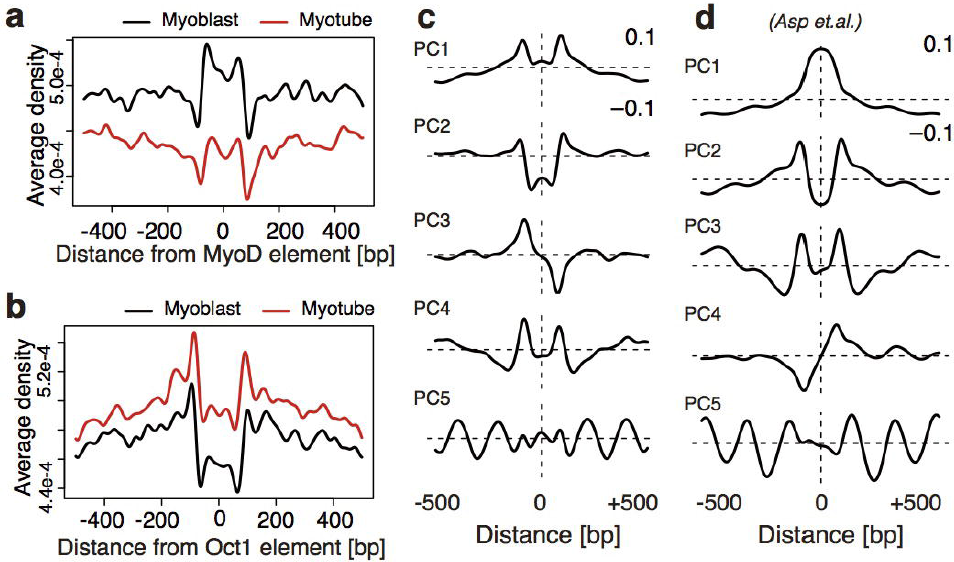
Five components extracted from PAND shapes around *cis*-elements. a. **(a, b)** PANDs of MyoD (a) and Oct (b) within 500 bp of their *cis*-elements. The X-axis denotes the distance (in bp) from the origin of the center of a *cis*-element, and the Y-axis denotes the nucleosome density averaged over all *cis*-elements.
b. **(d, d)** Principal components of the 258 PANDs. (c) The 1st to 5th principal components (top to bottom) were ordered by the contribution ratio and contributed approximately 80% of the cumulative contribution ratio. (d) The principal components of the ChIP-Seq input data from Asp et al (Asp et al. 2011).

Next, we attempted to extract common NP patterns across corrected PAND shapes using principal component (PC) analysis, since the PANDs were the result of averaging individual NPs at each locus of *cis*-elements (Fig. 2a,b and Supplementary Fig. 2). Figure 2c shows the top five PCs extracted from the analysis. Similar PCs were also extracted from MNase-treated data of C2C12 cells by Asp et al. (Fig. 2d) (Asp et al. 2011). The results show reproducibility of the components and also that the five components are invariable NP formations. The cumulative contribution ratio of the second PC was approximately 60%, showing that the majority of the 258 shapes can be explained by a combination of PC1 and PC2. Changes in PC1 and PC2 for the 258 *cis*-elements were observed in C2C12 cell differentiation, indicating NP alterations from the myoblast to myotube (Supplementary Fig. 3a). Therefore, PANDs could be used to extract NP dynamics.

**Figure 3.**
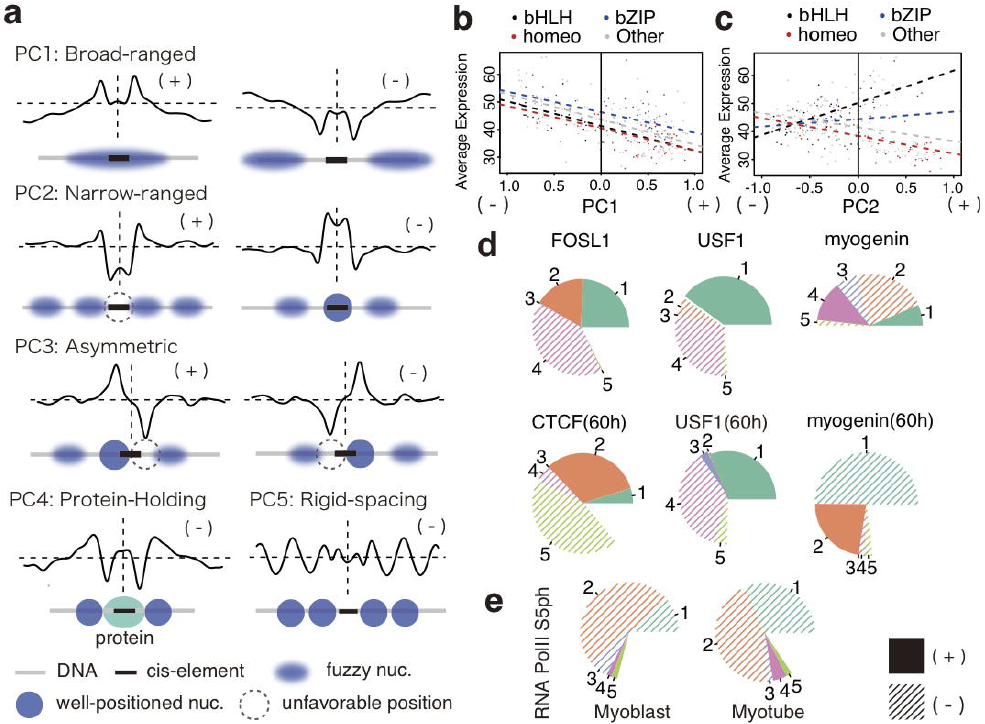
Functional profile of the five components. a. **(a)** Functional summary of the five components. (+)/(-) indicate positive/negative PCs in Figure 2c. The sign alternation results in a horizontal flipping of the component’s shape. Blue circles indicate well-positioned nucleosomes; blurred ovals indicate fuzzy positioned nucleosomes; and dotted circles indicate unfavorable positions of nucleosomes relative to *cis*-elements (black bar). The grey and black bars indicate DNA and *cis*-elements, respectively.
b. **(b, c)** Scatter plots of PC1/PC2 scores vs. average gene expression levels in myoblasts. The X-axis shows the PC1 (a) and PC2 (b) scores of cis-elements, and the Y-axis shows average expression levels of the genes covered by *cis*-elements. The regression lines of four groups (bHLH, bZIP, homeo, and Other) of cis-elements are shown as coloured dotted lines. The expression data (mRNA-Seq) were obtained from DRA000457.
c. **(d)** Functional profile of ENCODE TFBS data in mouse C2C12 cells. The content of each component is presented as a pie chart where the composition ratios (%) of five components are indicated. Numbers correspond to PC 1-5. Composition ratios from components other than PC1-5 are shown as blank regions.
d. **(e)** Functional profiling of RNAP2-Ser5ph in the myoblast/myotube shown in (c).

### Functional correspondences of five NP patterns

To interpret PCs as chromatin function, we further addressed the functional correspondence of the five PCs below.

#### *Broad-ranged NP* (PC1)

PC1 is characterized by a broad peak (+; in the case of positive PC1 score)/valley (-; negative score) like shape (Fig. 3a). This component reflects stochastic positioning rather than deterministic positioning and results in a relative anchor-point with NP at the center of PC1(+). PC1(-) corresponds to the NDR, which is a transcriptionally active domain. Finally, the average expression level of genes within 2 kbp of each *cis*-element were negatively correlated with the PC1 score (Fig. 3b, the Pearson’s correlation coefficient is −0.532), i.e. a positive PC1 tended to be associated with transcriptional repression, and a negative PC1 tended to be associated with transcriptional activation. Details of the gene sets are shown in Supplementary Figure 3b.

#### *Narrow-ranged NP* (PC2)

PC2 has an almost flat signal at base pairs far from the center, but a concave(+)/convex(-) shape near it (Fig. 3a). The shape suggests local accessibility within one nucleosome width that exposes or protects the DNA-binding sequence. PC2 had small negative correlation with the expression level in total, but a relatively large positive correlation in the bHLH group (Fig. 3c; Pearson’s correlation coefficients of each group are bHLH: 0.569, bZIP: 0.209, Homeo: −0.501, Other: −0.261 and −0.266 in total). This result indicates that transcription activation by the local/point arrangement of the nucleosome depends on specific TF types.

#### *Asymmetric NP* (PC3)

PC3 explains some of the asymmetrical shape of the PANDs (Fig. 3a). Such a shape was frequently found in the *cis*-elements of nuclear receptors (Supplementary Fig. 2) and could be explained by NP asymmetry and structural polarity.

#### *Protein-holding NP* (PC4)

PC4 is associated with the shape in which nucleosomes are stably fixed at positions greater than the ordinary nucleosome repeat length (186 bp) from the binding site (Fig. 3a). TF profiles of the five components revealed that PC4(-) makes a higher contribution in the ChIP-Seq data than in the 258 *cis*-elements (Supplementary Tables 1a and 1b), suggesting ChIP-Seq reports the actual binding site, not the *cis*-elements, which contain information on the bound and unbound state of the TF. The results suggest that PC4(-) includes a factor-holding state between nucleosomes. PC4(+) was not considered since the most of PC4 scores is negative (-) in PANDs of *cis*-elements.

#### *Rigid-spacing* NP (PC5)

PC5 exhibited NP that had equal intervals on both sides of a TFBS (Fig. 3a). A large PC5(-) composition ratio was seen at the CTCF binding site (Fig. 3d), suggesting a boundary formation on the genome (Supplementary Tables 1a).

### TF function explained by the composition of five NP patterns

To evaluate whether our approach can unveil TF function in transcriptional regulation, PANDs were generated and used to profile various TFBSs in C2C12 cells from the ENCODE project (Fig. 3d) (Stamatoyannopoulos et al. 2012). The profile of FOSL1 showed large PC1(+) and PC4(-) in the myoblast (unlabeled; 0 h), and USF1 showed a similar and consistent profile in both myoblasts and myotubes (60 h). These results suggest a function of stably-bound and constitutive transcriptional suppression by FOSL1 and USF1 during differentiation (Pognonec et al. 1997). Additionally, FOSL1 has a bZIP motif and can activate transcription (Pognonec et al. 1997); thus coupled with the AP-1 family, PC2(+) in FOSL1 indicates the other side of FOSL1 is involved in transcription activation (see bZIP group in Fig. 3c). On the other hand, myogenin showed changes in its composition ratio during differentiation (Fig. 3d). The PAND of myogenin at 0 h was not well characterized by the five components (i.e. half of the compositions were blank) because myogenin is not expressed in the myoblast, which reflects the fact that myogenin at 0 h did not contribute to transcription activation. However, PANDs had large composition rates for PC1(-) and PC2(+) at 60 h, which suggests an active transcription state. These results are consistent with a reported function of myogenin in transcription (Ohkawa et al. 2007).

### ChIP-Seq could predict TF function when combined with fixed MNase-Seq

MNase is sometimes utilized to prepare input samples for ChIP-Seq (Zhang et al. 2008). Accordingly, we attempted to perform ChIP-Seq and fixed MNase-Seq simultaneously to explore TF involvement in transcription. We performed ChIP-Seq on the RNAP2 phosphorylated carboxyl-terminal domain (CTD) serine-5 (RNAP2-S5ph) using MNase digestion following fixation. After identifying RNAP2-S5ph loading sites on the genome, PANDs around RNAP2-S5ph were profiled by the composition of the five PCs. The shapes of the PANDs for myoblasts and myotubes were mostly due to PC1 and PC2, with the PC1(-) content being much higher in the myotube stage than in the myoblast (Fig. 3e). Because our RNAP2-S5ph antibody recognizes both the poised state (S5ph) and the active transcribing state (S5ph + S2ph) (Brookes and Pombo 2009; Odawara et al. 2011), the increase in PC1(-) suggests an increase in transcriptional activation around the RNAP2-S5ph binding sites, which is consistent with a previous finding that showed increased transcription of activated genes during skeletal muscle differentiation (Harada et al. 2012).

### Co-positioning nucleosome detection for five NP patterns

Evaluating the co-existence of nucleosome pairs confirmed that the five components represent NP at individual gene loci (Fig. 4). The shapes of the five components reflect NP at a single locus. However, it is difficult to reconstruct the actual NP because a PAND just represents the average frequency of single nucleosome detection. Even if a triple-peaked PAND is obtained, nucleosomes at the peak do not necessarily appear at the same time for each gene locus (Fig. 4a). Therefore we confirmed whether a single PAND that shows more than two peaks reflects NP at an individual gene locus. The degree of co-coexistence was visualized by the evaluation of cosine-similarity of nucleosome pair detection at all possible combination of positions. This visualization extracts pairs of positions where nucleosomes appear frequently and commonly around anchor-points, so that individual NP for a gene locus can be confirmed. Thus, co-occurrence evaluation proved the existence of NP as predicted from the five PCs at an individual gene locus (Fig. 4b-f).

**Figure 4.**
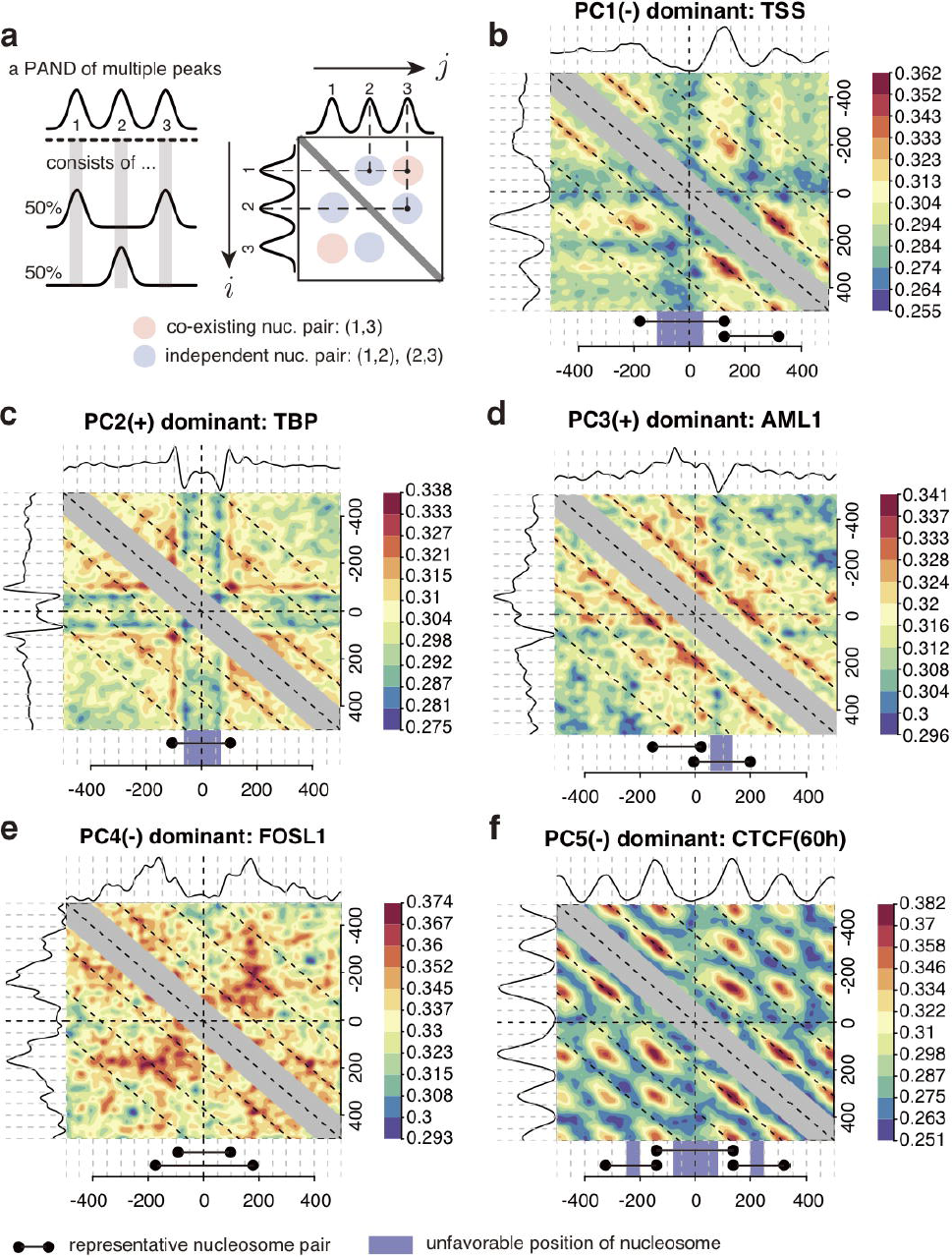
Frequently appearing nucleosome pairs. **(a)** Scheme of the nucleosome pair extraction method. A PAND with three peaks: 1, 2 and 3, which indicate three nucleosomes, is shown. However, the PAND is not sufficient for analyzing nucleosomes that appear at the same locus (a, left). Therefore the degree of co-existence at two arbitrary points around an anchor-point was visualized (a, right). The pairs (1,2) and (2,3) indicate nucleosome pairs that appear independently, while the pair (1,3) is a co-existing nucleosome pair at a single locus. **(b-f)** Examples of NPs based on PC1-5. The respective PAND shape is located at the top and left of each matrix. The color-bars indicate the cosine similarity of a position pair in a matrix. Representative NPs are shown at the bottom of the matrices. **(b)** The TSS has a PC1(-) dominant shape with a high co-occurrence of (+1,+2) and (-1,+1) nucleosomes. **(c)** TBP has a PC2(+) dominant shape that is consistent with nucleosomes tending not to position at the center of the *cis*-element. **(d)** AML1 has a PC3(+) dominant shape, which indicates nucleosomes are well-positioned near the *cis*-element, while an unfavorable position exists approximately +100 bp from the cis-element. **(e)** Fosl1 has a PC4(-) dominant shape that indicates NP where two nucleosomes are positioned approximately 300 bp from the binding site in opposite directions. Since the PAND of Fosl1 also contains PC2(+), two exclusively existing NP of PC4(-) and PC2(+) were observed. **(f)** The CTCF binding site has a PC5(-) dominant shape.

## DISCUSSION

Here we proposed a framework that applies single fixed MNase-Seq data to acquire profiles of TF function. We found specific NP patterns that indicate transcription activation and inhibition. Our results suggest that five NP patterns indicate the existence of a structural code defined on the chromatin and describes the transcription state of the TFs. Similar chemical modifications of histones or RNAP2 have been reported as histone or CTD codes that can describe the transcription regulation state (Jenuwein and Allis 2001; Buratowski 2003; Segal et al. 2006). These codes have the potential to act as detailed spatial descriptions of the chromatin structure.

Our method assumes that most nucleosome-wrapping DNAs produce mono-nucleosome sized DNA fragments. However, some fragments, such as small histone alternative proteins, non-nucleosomal proteins or poly-nucleosome may produce variable fragment sizes (Henikoff et al. 2011). Furthermore, MNase-Seq data include various types of nucleosomes with non-canonical histone variants, such as H2A.Z and H3.3. Further understanding of how complex NP conformations contribute to transcription regulation requires more structural characterization of the NP such as analysis of the histone variants associated with the nucleosome, as this would provide more information on the flexibility of the DNA-nucleosome interaction, co-localizations of histone-variants on the genome, and contribution to histone-modifications and gene expression.

Another limitation of our method is that the evaluation depends on averaging: NP evaluation was not focused to individual NP at single loci and/or in single cells. Recently, a computational reconstruction method of single NPs from observed MNase-Seq signals in single gene loci was proposed (Schöpflin et al. 2013). Although this method enabled NP estimation at each single site, a large number of reads is still required to recapitulate single NP level. In our case, to confirm NPs formed by poly-nucleosomes at single loci, we evaluated the co-existences of nucleosomes in a statistical aspect (Fig. 4). Further approaches for revealing the spatial arrangements of poly-nucleosomes (two or more) by MNase-Seq should allow for direct comparison between our results and biochemistry or microscopy studies.

We demonstrated our approach reproduces previous findings (Fig. 2d) and can be applied to the ENCODE data of C2C12 cells, which suggests its applicability to other ChIP-Seq data for the prediction of TF function in transcription regulation. Although this study focused on TFs, the same method could be applied to specific histone modifications or other epigenetic markers on the genome. To better understand the function of transcription-independent NP patterns, such as narrow-ranged, asymmetric and protein-holding, however, other functional predictions and understanding of other NP patterns are needed for study of higher-order chromatin structures.

## METHODS

### Cells

C2C12 cells were cultured in Dulbecco’s modified Eagle’s medium (DMEM) supplemented with 20% fetal bovine serum. Cells examined under growth conditions (myoblast) were harvested at 60-70% confluency. Differentiated samples (myotube) were transferred to DMEM containing 2% horse serum upon reaching confluence and harvested 48 h later.

### MNase-digestion, chromatin immunoprecipitation and deep sequencing

Cells were fixed with 0.5% formaldehyde for 5 min at room temperature and then blocked by 150 mM glycine, pH 7.0. After sonication three times (5 sec each time), the cells were digested with MNase for 40 min at 37 °C. The rest of the procedure is described by Odawara et al. (Odawara et al. 2011). ChIPed DNA was sequenced (Illumina HiSeq 2000; Illumina K.K.), and the obtained reads were mapped to the mouse genome (mm9) using Bowtie2 (version 2.1.0) software with default parameter settings. Multi-hit reads were discarded. The binding sites of RNAP2-S5ph were obtained with MACS (Zhang et al. 2008) (version 2.0.10.20130528) using the options “*—gsize mm —nomodel —extsize (average ChIPed DNA fragment size, 170 for myoblast and 159 for myotube) —broad —to-large —pvalue 1e-3*”.

### Density estimation of the nucleosomes

Nucleosome density was estimated as follows:

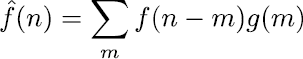

where *f*(*x*) is the normalized count of the nucleosome mid-point at genomic coordinate *x* (in bp) using the reads per million normalization (Mortazavi et al. 2008). The estimated density of a nucleosome mid-point, 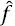, was calculated by a linear convolution of *f* and the Gaussian kernel, *g*,

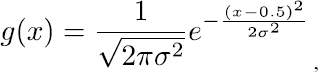

where σ is the bandwidth parameter. We calculated the convolution over the range *m*∈ [*−4σ*, 4σ], because *g*(*x*) becomes sufficiently small to ignore when | *x*| > 4σ. The term *x* – 0.5 is a continuity correction.

The parameter σ can represent the statistical indeterminacy of the mid-point of the DNA fragment (i.e. center of nucleosomal DNA) that was estimated from a single-end read. The optimal setting of σ was determined experimentally by measuring the DNA fragment size. We obtained an average fragment size of 164 bp +/−12 bp (standard deviation of fragment size) from an MNase-digested sample of the myoblast. This measurement provides statistical evidence that a nucleosome mid-point positions 82 (half size of the average) +/−12 bp (2 standard deviations) from the 5' end of a mapped read; the probability was 95.45%. We therefore used σ = 12 bp as the bandwidth parameter, as it is sufficiently large to cover each center of all nucleosomal DNA.

### Genomic region definitions of 258 *cis*-elements

258 *cis*-element-containing region definitions were acquired from the University of California Santa Cruz (UCSC) TFBS Conserved Track (*tfbsConsSites*). The track estimates *cis*-element-containing regions from the Transfac Matrix Database (v7.0) of the human genome (hg19). The region definition was converted to the mouse genome (mm9) using UCSC’s *liftOver* tool.

### ENCODE ChIP-Seq TFBS data

TFBSs by ChIP-seq from ENCODE/Caltech (GSE36024) were used to profile TF function with PANDs. The calculated TFBSs from the data, named *narrowPeak*, were obtained from UCSC: http://hgdownload.cse.ucsc.edu/goldenpath/mm9/encodeDCC/wgEncodeCaltechTfbs.

### Principal component analysis of average density profiles

A data matrix, **X**, of 1001 × 516 elements was generated. Columns contain PANDs for 258 types of *cis*-elements ± 500 bp in the myoblast/myotube. Each column was normalized to average = 0 and Euclid norm = 1. This normalization was used to compute the composition ratio of each principal component. The eigenvectors of the covariance matrix, 1/(516-1) **XX**^*t*^, were used as principal components. The cumulative contribution ratio was computed from the eigenvalues.

### Extraction of frequently appearing nucleosome pairs

The co-existence of nucleosomes detected simultaneously at two arbitrary points around an anchor-point was computed using smoothed density profiles at individual sites. We defined **A** as a matrix that holds all combinations of two arbitrary points within +/−1 kbp of the anchor-point (e.g. TSS or center of *cis*-element). Its elements are cosine similarities for all combinations of *i, j* ∈ [−1000,1000], where the cosine similarity is defined as

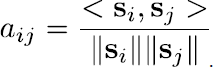

The vectors *s*_*i*_ and *s*_*j*_ are the *i*-th and *j*-th columns of matrix **S**, an *M* × *N* matrix, where *M* is the total number of individual sites and *N* is the range (in bp) of the nucleosome density. We used *N* = 1001, i.e. +/− 1 kbp from the anchor-point. The notation ||.|| is a Euclid norm (root sum squared), and the function (·,·) is the dot product of vectors.

## END NOTES

### Data Access

Fixed MNase-Seq data and ChIP-Seq data of RNAP2-S5ph in myoblast/myotube cells were submitted to DDBJ Sequence Read Archive with the accession number: DRA001262.

## Acknowledgments

We thank H. Kurumizaka, H. Kimura, Y. Arimura for insightful discussions; T. Ichinose, M. Kato, N. Urasaki for technical support; A. Harada, P. Karagiannis, A. N. Imbalzano for reading the manuscript; and Research Institute for Information Technology, Kyushu University (tatara) and National Institute of Genetics (NIG) for providing the high-performance computing resources. This work was supported by the Core Research for Evolutional Science and Technology (CREST), JSPS KAKENHI Grant Number 23310134, 25116010, 25132709, 25118518.

## Author Contributions

KM performed the experiments and writing the manuscript and YO wrote the manuscript and designed the experiments.

